# Klotho deficiency promotes skeletal muscle weakness and is associated with impaired motor unit connectivity

**DOI:** 10.1101/2025.06.11.659129

**Authors:** Linda A. Bean, Connor Thomas, Juan F. Villa, Alexander J. Fitt, Areli Jannes S. Javier, Akanksha Agrawal, Hanna Whitney, Guilherme Nascimento Dos Santos, Kenneth E. White, Joshua R. Huot, Steven S. Welc

**Affiliations:** Department of Anatomy, Cell Biology & Physiology Indiana University School of Medicine Indianapolis, IN, USA; School of Medicine Universidad CES Medellin, Colombia; Indiana Center for Musculoskeletal Health Indiana University School of Medicine Indianapolis, IN, USA; Department of Medical and Molecular Genetics Indiana University School of Medicine Indianapolis, IN, USA

**Keywords:** Klotho, skeletal muscle, wasting, motor unit

## Abstract

Muscle wasting and weakness are important clinical problems that impact quality of life and health span by restricting mobility and independence, and by increasing the risk for physical disability. The molecular basis for this has not been fully determined. Klotho expression is downregulated in conditions associated with muscle wasting, including aging, chronic kidney disease, and myopathy. The objective of this study was to investigate a mechanistic role for Klotho in regulating muscle wasting and weakness. Body weight, lean mass, muscle mass, and myofiber caliber were reduced in Klotho-deficient mice. In the tibialis anterior muscle of Klotho null mice, type IIa myofibers were resistant to changes in size, and muscle composition differed with a higher concentration of type IIb fibers to the detriment of type IIx fibers. Glycolytic enzymatic activity also increased. The composition of the soleus muscle was unaffected and myofiber caliber was reduced comparably in type I, IIa, and IIx fibers. Muscle contractile function declined in Klotho-deficient mice, as evidenced by reduced absolute twitch and torque, and decreased rates of contraction and relaxation. RNA-sequencing analysis identified increased transcriptional expression of synaptic and fetal sarcomeric genes, which prompted us to test effects on muscle innervation. Klotho-deficiency induced morphological remodeling of the neuromuscular junction, myofiber denervation, and a functional loss of motor units. Loss of motor units correlated with absolute torque. Collectively, our findings have uncovered a novel mechanism through which Klotho-deficiency leads to alterations to the muscle synapse affecting motor unit connectivity that likely influences muscle wasting and weakness.

**Key points summary:**

- Maintaining skeletal muscle mass and function is critical to preserve physical capacity and independence. Clinical observations implicate longevity factor ⍺Klotho as a key regulator of muscle mass and weakness. Low Klotho levels are reported to correlate with muscle weakness and frailty.
- Using Klotho null mice, our study shows that Klotho-deficiency promotes skeletal muscle weakness and impaired motor unit connectivity.
- RNA-sequencing analysis identified altered expression of sarcomeric and synaptic genes suggesting changes to the muscle synapse in Klotho-deficient mice.
- Histopathological analyses revealed Klotho-deficiency is associated with reduced myofiber caliber, altered muscle composition, and increased prevalence of NCAM+ denervated fibers. Imaging of the NMJ further showed morphological changes and reduced area of synaptic contact.
- Overall, our findings show that Klotho regulates the structure and function of the NMJ affecting motor unit connectivity which may have an important role in the pathogenesis of muscle wasting and weakness.

## INTRODUCTION

The long-term ability of individuals to preserve their physical capacity and independence largely depends on maintaining skeletal muscle mass and function. Muscle wasting facilitates weakness and an overall decline in physical function, increasing the risk of falls and injury that lead to disability (Wolfson *et al*., 1995; Tinetti & Williams, 1997; Liu *et al*., 2014; Mayer *et al*., 2020). Furthermore, systemic metabolic adaptations occurring with changes in skeletal muscle function may act as a disease modifier affecting pathophysiology and prognosis (Baskin *et al*., 2015). Muscle wasting and persistent weakness impact quality of life and limit the ability of individuals to perform normal daily activities, to work to earn a living, or to return to work after critical illness, owing to considerable societal burden and health care expenditure (Hermans & Van den Berghe, 2015; Pinedo-Villanueva *et al*., 2019). With aging being a well-known contributing factor to loss of muscle mass, the costs of muscle weakness and its associated complications will continue to rise dramatically as the aging population rapidly expands in size (Kalyani *et al*., 2014). While there is considerable evidence to support that muscle mass and function are strong prognostic indicators in various medical conditions, the underlying molecular regulators of muscle mass and function have yet to be fully established.

Mutations to the ‘aging suppressor’ Klotho gene results in a shortened lifespan and changes to numerous tissues resembling pre-mature aging (Kuro-o *et al*., 1997). The Klotho gene encodes a transmembrane protein with a large extracellular component. Klotho expression is restricted to several tissues, including skeletal muscle. However, its function is best known where it is abundantly expressed, in the kidney, acting as a co-receptor for fibroblast growth factor 23 to regulate phosphate and mineral homeostasis (Urakawa *et al*., 2006). The extracellular fraction can be enzymatically cleaved and released as a soluble factor with discrete functions modifying insulin/IGF-1, TGFβ, NF-κB, Wnt, and NRF2 signaling (Kuro-o, 2019).

Klotho may function as a regulator of muscle mass and weakness. Both circulating Klotho levels and muscle mass begin to decline after 40-years-old (Xiao *et al*., 2004). Low Klotho correlates with frailty and increased occurrences of falls in nursing home residents (Shardell *et al*., 2019; Sanz *et al*., 2021; Veronesi *et al*., 2021). Plasma Klotho levels are negatively associated with daily living disability and positively associated with lean mass (Crasto *et al*., 2012; Shardell *et al*., 2020). Low Klotho levels also correlate with poor muscle strength and early mortality (Semba *et al*., 2012).

In the present study, we tested the effects of Klotho-deficiency on muscle mass and function. We assayed for changes in composition of fast-twitch tibialis anterior (TA) and slow-twitch soleus muscles, and distinguished morphology by fiber-type. Klotho-deficiency resulted in a shift from type IIx toward a higher proportion of type IIb fibers in TA muscles, which was paralleled by increased activity of the glycolytic enzyme, ⍺-glycerolphosphate dehydrogenase (GPDH). We identified transcriptional reprogramming associated with altered expression of sarcomeric and synaptic genes. We then extended our hypothesis beyond muscle-level changes to test if Klotho-deficiency affects motor unit connectivity, a determinant of muscle function. We used an established motor unit number estimation (MUNE) technique to evaluate number of motor neurons functionally connected to muscle. Histological assessment further demonstrated enhanced expression of denervation marker neural cell adhesion molecule (NCAM), reduced area of synaptic contact, and neuromuscular junction (NMJ) morphological alterations. Taken together, these results support an important role for Klotho in regulation muscle mass, composition, contractile and neuromuscular function.

## METHODS

### Ethical approval

All animal experiments were performed according to approved Institutional Animal Care and Use Committee protocols at Indiana University School of Medicine conforming with federal ethical regulations, AAALAC standards for animal testing and research, and with the 1964 Declaration of Helsinki and its later amendments.

### Mice

Klotho^+/-^ mice were obtained from Taconic Biosciences (#TF0361, Germantown, NY) and bred to generate wild-type (WT) and homozygous (KL^-/-^) mice. Genomic DNA was extracted from tail snips or ear punches using hot sodium hydroxide and tris (Truett *et al*., 2000). Genotype was determined by PCR reaction (Promega #M8296) and primers for Klotho mutant (F: 5’-ATGCTCCAGACATTCTCAGC-3’ and R: 5’-GCAGCGCATCGCCTTCTATC-3’) and control products (F: 5’-GATGGGGTCGACGTCA-3’ and R: 5’-TAAAGGAGGAAAGCCATTGTC-3’) separated by agarose gel electrophoresis. Weaned mice were housed in static cages containing environmental enrichment in a specific pathogen free facility maintained at 21–22°C with a 12-hour light/dark cycle with chow diet (2018SX, Teklad, St. Louis, MO) and water provided ad libitum. Body composition (EchoMRI™-700, Houston, TX), *in vivo* plantarflexion torque, and motor unit connectivity were assessed in male mice 45-50-days-old prior to effects on mortality (Nakatani *et al*., 2009). At time of euthanasia, muscles were harvested, weighed, and frozen in liquid nitrogen or embedded in OCT compound and frozen in liquid nitrogen-cooled isopentane.

### Immunofluorescence and Morphological Analysis

Frozen 10 μm transverse and 20 μm longitudinal muscle sections were blocked in PBS with 0.1% tween 20 containing 5-10% normal donkey serum. Sections were immunolabeled as performed previously (Javier *et al*., 2025). Antibodies used were laminin (1:200; #L9393) and NCAM (1:250; #AB5032) from MilliporeSigma (Burlington, MA); myosin heavy chain (MyHC) I (1.2 μg/ml; #BA-D5), IIa (1.3 μg/ml; #SC-71), IIb (1.9 μg/ml; #BF-F3), dystrophin (2 μg/ml; #Mandys8(8H11)), synaptic vesicles (0.54 μg/ml; #SV2), and neurofilament (0.46 μg/ml; #2H3) from Developmental Studies Hybridoma Bank (Iowa City, IA). Sections were then probed with fluorochrome-conjugated secondary antibodies anti-mouse IgG2b (#A21242), IgG1 (#A21121), IgM (#A21044), and anti-rabbit IgG (#A11046) from Invitrogen (Carlsbad, CA); or anti-mouse IgG (#715-585-151) and anti-rabbit IgG (#711-545-152) from Jackson ImmunoResearch (West Grove, PA). Acetylcholine receptors (AChR) were labeled with αbungarotoxin (1:500; #0007, Biotium, Fremont, CA) and nuclei with DAPI. Sections were imaged using a Zeiss AxioObserver 7 microscope as previously described (Earl *et al*., 2024).

Morphological analysis was performed by segmenting myofibers from single channel fluorescence images of laminin-stained sections using Cellpose (v2.2.3) (Stringer *et al*., 2021). Images from Cellpose segmentation were created in ImageJ (v2.1.0/1.53c) and Regions of Interest were eroded with a fixed number of pixels using the plugin LabelsToRois. The resulting ROIs were applied to the original multi-color image for quantification of myofiber cross-sectional area (CSA) and fiber-type. Myonuclear number and central nucleation were quantified from dystrophin labeled sections stained with DAPI. For quantification of NMJ morphology, images were acquired in a z-series at 0.5 µm intervals, raw stacks were deconvolved using the nearest-neighbor algorithm and then collapsed using the extended depth-of-focus module. Individual NMJ inset images were duped out of the full image, then split into separate channels, red and green channels were merged to create a 2-color 2-channel hyperstack for processing using an ImageJ-based macro NMJ-morph (Minty *et al*., 2020).

### Histochemistry

To assess oxidative capacity, succinate dehydrogenase (SDH) activity was assayed by incubating frozen sections in 0.1M phosphate buffer, pH=7.0, with 1.5 mM nitrotetrazolium blue chloride, 130 mM sodium succinate, 0.2 mM phenazine methosulfate, and 1 mM sodium azide (Blanco *et al*., 1988). GPDH activity was assayed as an in-direct measure of glycolytic activity, frozen sections were incubated in 0.1M phosphate buffer, pH=7.0, with 1.2 mM nitrotetrazolium blue chloride, 0.8 mM phenazine methosulfate, 9.3 mM glycerol phosphate disodium salt hydrate, and 1 mM sodium azide (Martin *et al*., 1985). Background staining was assessed in negative control slides by omitting enzyme substrates. Serial sections were used to determine enzyme activity in individual myofibers based on immunofluorescence labeling for MyHCs. Enzymatic activities of all myofibers on a whole section was performed by segmentation using Cellpose and quantification of optical density using ImageJ.

### Immunoblot

Protein extraction and immunoblot assays were performed as done previously (Javier *et al*., 2025). Muscle (50 µg) and kidney (5 µg) lysates were subjected to SDS-PAGE and proteins transferred onto nitrocellulose membranes. Equal protein loading was assessed using Ponceau S. After blocking membranes with 5% non-fat dry milk, membranes were incubated in anti-Klotho (1:500; #MABN1807, MilliporeSigma) in blocking solution and probed with HRP-conjugated anti-rat (#7077, Cell Signaling Technologies, Danvers, MA) antibody. Bands were visualized using SuperSignal West Femto chemiluminescence substrate (ThermoScientific) and captured using a Chemidoc Imager (BioRad, Hercules, CA).

### Serum Analyses

Creatine kinase (CK) activity (#MAK116, MilliporeSigma) and phosphate levels (#ab65622, Abcam, Cambridge, MA) were measured in serum using commercially available assays. Absorbance was read using a SpectraMax M2 microplate reader (Molecular Probes, Eugene, OR) and concentrations calculated using regression analysis.

### RNA Isolation, Quantitative Real-time PCR (qRT-PCR), RNA-sequencing, and Bioinformatic Analysis

RNA isolation and qRT-PCR assays were completed as performed previously (Earl *et al*., 2024). Relative expression of selected transcripts (Table 1) was normalized by geometric averaging the Cq values of 2 references genes. The expression of each gene in control samples was set to one and other expression values were then scaled to that value. RNA quality assessment, library construction, and mRNA sequencing was performed at the Indiana University School of Medicine Center for Medical Genomics. Briefly, 100 ng of RNA was used to generate libraries. Libraries were sequenced on a NovaSeq 6000 (Illumina, San Diego, CA) generating approximately 40M reads per library. Quality control of raw sequence data was performed with FastQC (v0.11.5) and sequencing reads mapped to the mm10 mouse reference genome using STAR (v2.7.10a). Low quality mapped reads were excluded, featureCounts was used to quantify gene level expression, and differential expression analysis was performed with edgeR. Gene Set Enrichment Analysis (GSEA) with Gene Ontology (GO) and Kyoto Encyclopedia of Genes and Genomes (KEGG) pathway functional enrichment analyses were performed using clusterProfiler.

**Table 1:**
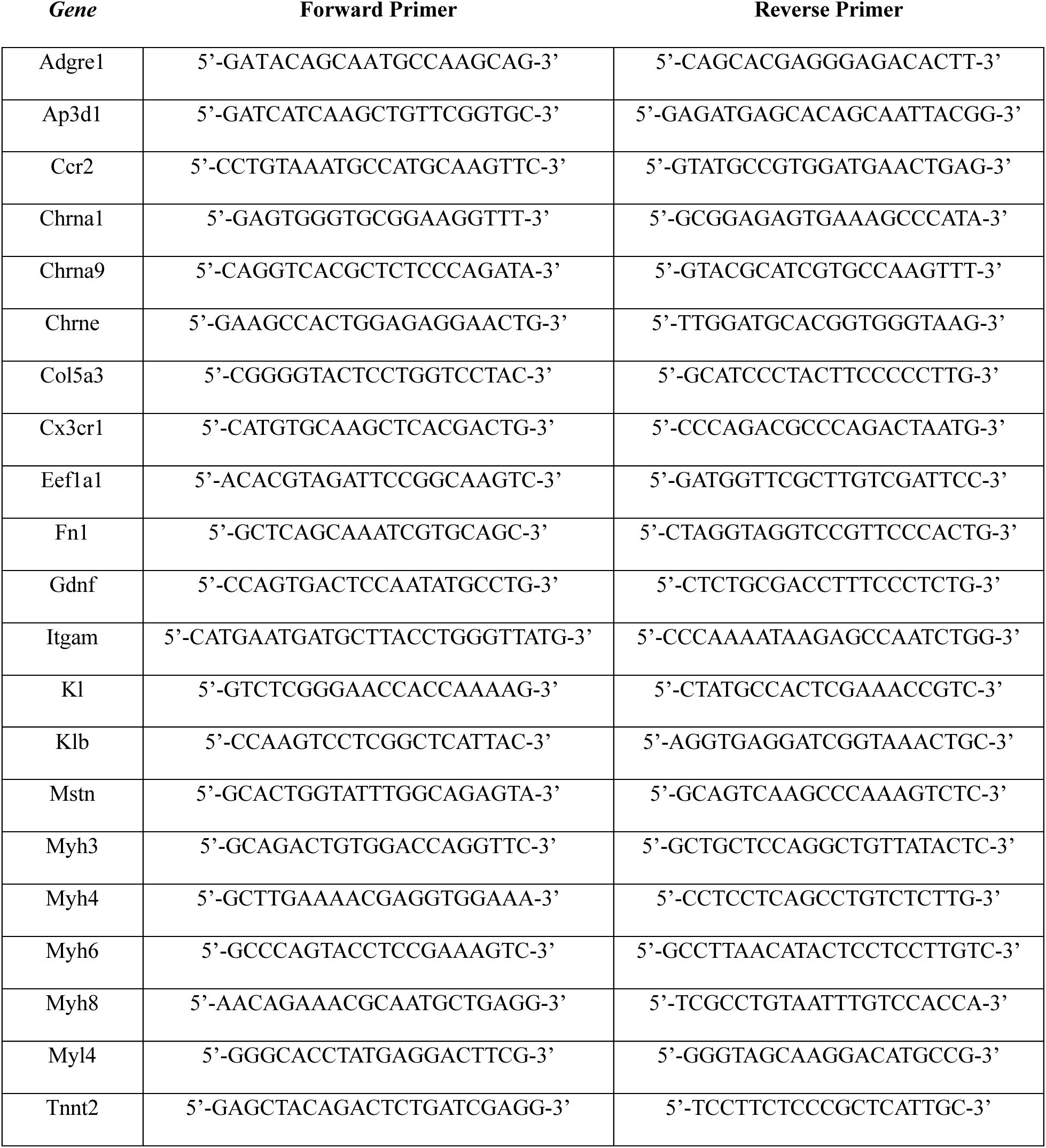
qRTPCR Primers.

### In Vivo Plantarflexion Torque Assessment

Mice underwent *in vivo* plantarflexion torque assessment to assess contractility of the triceps surae (Aurora Scientific, Aurora, ON). Briefly, mice were anesthetized with 2% isoflurane inhalation and the right hindfoot was taped to the force transducer positioned at a 90° angle to the tibia, and the limb was fixed by knee-clamp, avoiding compression of the fibular nerve. Two disposable monopolar electrodes (Natus Neurology, Middleton, WI) were inserted to stimulate the tibial nerve. Maximum twitch torque was first determined using supramaximal stimulations (0.2 ms square wave pulse). Peak plantarflexion torque was then assessed following a supramaximal square wave stimulation (0.2 ms) delivered at a frequency of 125 Hz. Peak tetanic torque, maximum rate of force contraction and relaxation were analyzed using Dynamic Muscle Analysis software.

### In Vivo Electrophysiology

Electrophysiological assessment was used to evaluate compound muscle action potential (CMAP) and MUNE as performed previously (Huot *et al*., 2021). The sciatic nerve of the left limb was stimulated with two 28-gauge needle electrodes (Natus Neurology), a duo shielded ring electrode was used for recording, and a ground electrode was placed on the animal’s tail. Baseline-to-peak and peak-to-peak CMAP responses were recorded utilizing supramaximal stimulations (constant current intensity: <10 mA; pulse duration: 0.1 ms). Single motor unit potential (SMUP) size was determined using an incremental stimulation technique. Incremental responses were obtained by submaximal stimulation of the sciatic nerve until stable, minimal all-or-none response occurred. Ten successive SMUP increments were recorded and averaged. MUNE was calculated by: CMAP amplitude (peak-to-peak)/average SMUP (peak-to-peak).

### Statistics

All data are presented as mean ± s.e.m. Statistical significance was calculated by unpaired two-tailed Student’s *t*-test or two-way ANOVA with Holm-Sidak or Tukey’s multiple comparison test using GraphPad Prism (v10). Differences with P<0.05 were considered statistically significant.

## RESULTS

### Genetic ablation of Klotho reduces lean mass, fat mass, and muscle mass

Previous studies have shown that reductions in systemic soluble Klotho concentrations are suppressed with aging and associated with muscle atrophy (Kuro-o *et al*., 1997; Xiao *et al*., 2004). Klotho levels are also reduced locally in skeletal muscles with aging, acutely in response to muscle injury, and chronically after the onset of inflammation in dystrophic muscles.(Wehling-Henricks *et al*., 2016; Wehling-Henricks *et al*., 2018; Welc *et al*., 2020) The first objective of this study was to examine the effects of Klotho-deficiency on skeletal muscle wasting. Mouse genotypes were determined by PCR (Fig. 1A) and immunoblots confirmed the absence of Klotho expression in KL^-/-^ kidney and muscle lysates (Fig. 1B). Normal muscles expressed full-length (130-KDa) Klotho protein. qRT-PCR also showed that Kl mRNA was not detectable in KL^-/-^ muscles, whereas Klb expression was normal (Fig. 1C). As expected, serum phosphate levels were elevated in KL^-/-^ mice (WT: 10.6±0.8 *vs.* KL^-/-^: 14.8±1.2 mg/dL, P<0.05).

**Figure 1:**
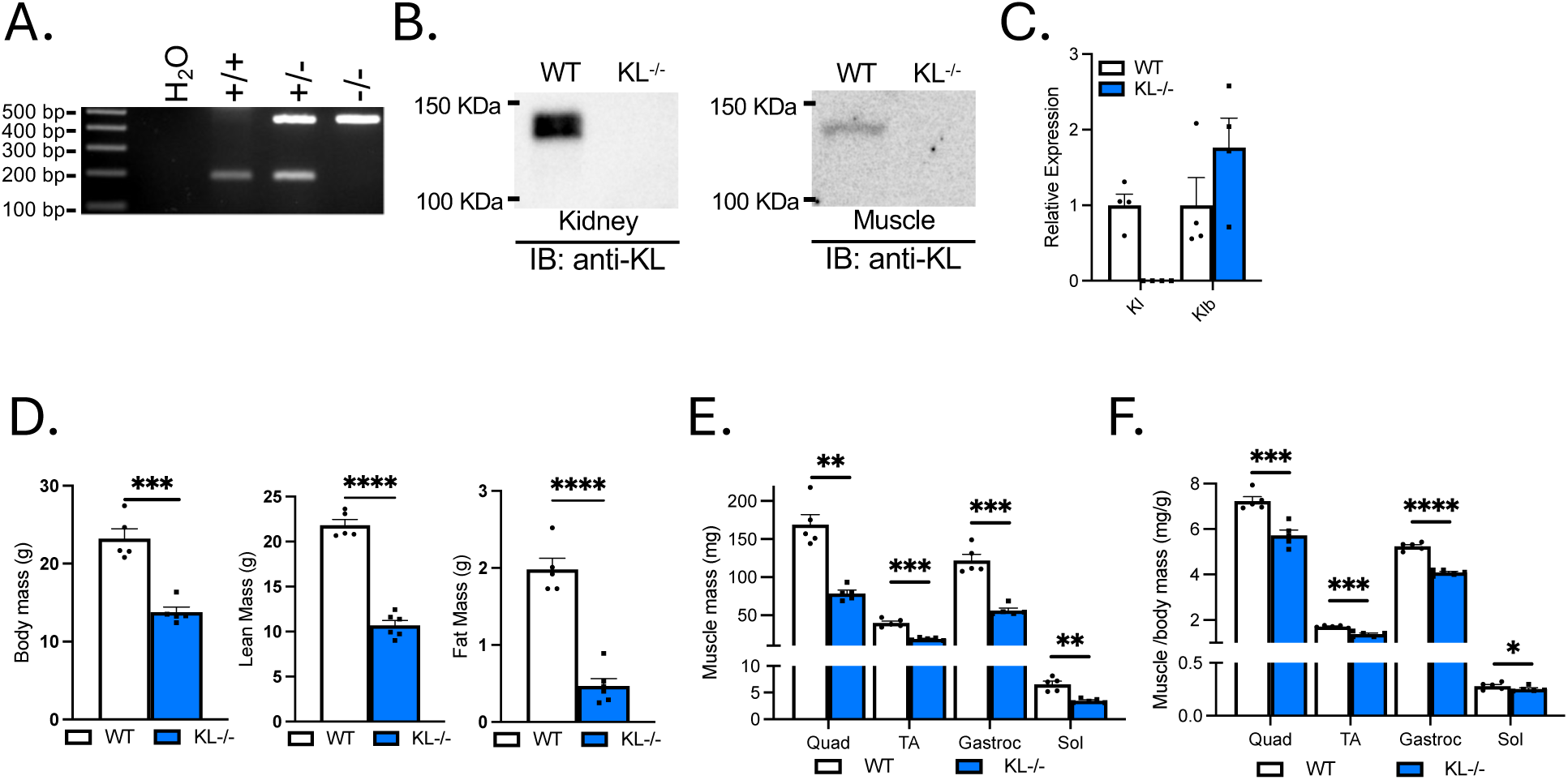
Klotho-deficient mice have reduced body mass, lean mass, fat mass, and muscle mass. (A) Representative image of wild-type (186 bp) and mutant (426 bp) PCR products from wild-type, heterozygous, and homozygous mice distinguish genotype. (B) Immunoblot confirms no detectable Klotho expression in kidney or muscle lysates of KL^-/-^ mice. (C) qRT-PCR assays show *Kl* mRNA expression is not detectable and *Klb* mRNA expression was normal in KL^-/-^ muscle lysates. N = 4 per group. (D) Body mass and body composition was assessed by Echo-MRI showing decreased lean mass and fat mass in KL^-/-^ mice. Muscle mass (E) and muscle mass normalized to body mass (F) of quadriceps (quad), tibialis anterior (TA), gastrocnemius (gastroc), and soleus (sol) in wild-type and KL^-/-^ mice. N = 5-6 per group. Data are presented as mean ± SEM. All *P* values are based on two-tailed *t* test. \**P*<0.05, \*\**P*<0.01, \*\*\**P*<0.001, \*\*\*\**P*<0.0001 *versus* wild-type.

Klotho-deficiency resulted in a ∼40% decrease in body mass (P<0.001, Fig. 1D). Body composition assessment revealed a reduction in lean (−51%, P<0.0001) and fat mass (−76%, P<0.0001, Fig. 1D). The gross wet mass of quadriceps (−53%, P<0.01), TA (−52%, P<0.001), gastrocnemius (−54%, P<0.001), and soleus muscles (−48%, P<0.01) were reduced in KL^-/-^ mice (Fig. 1E). Muscle mass to body mass ratio were also reduced in quadriceps (−21%, P<0.001), TA (−19%, P<0.001), gastrocnemius (−22%, P<0.0001), and soleus muscles (−11%, P<0.05) indicating that effects on muscle mass exceeded allometric scaling (Fig. 1F).

### Klotho-deficiency reduces myofiber size and affects muscle composition in fast-twitch TA muscles

Myofiber CSA was assayed to determine if reductions in muscle mass are due to differences in size. Mean myofiber size was reduced in TA muscles of KL^-/-^ mice (−41%, P<0.01) marked by an increase in the proportion of small fibers (≤999 µm^2^) and a reduction in the percentage of large fibers (1500-2499 µm^2^; Fig. 2A). Next, we evaluated changes in CSA by fiber-type on sections immunolabeled with antibodies to MyHC type I, IIa, and IIb. Type IIx fibers were identified by the absence of staining. The CSA of type IIx (−41%, P<0.05) and IIb fibers (−47%, P<0.0001) were reduced (Fig. 2B, D). A non-significant reduction in type IIa fiber size (−26%, P<0.10) suggests fiber-type related resistance to the effects of Klotho-deficiency. Muscle composition was altered with increased frequency of type IIb fibers (WT: 55.7% *vs.* KL^-/-^: 70.1%, P<0.01) at the expense of type IIx fibers (WT: 29.6% *vs.* KL^-/-^: 14.6%, P<0.01, Fig. 2C-D).

**Figure 2:**
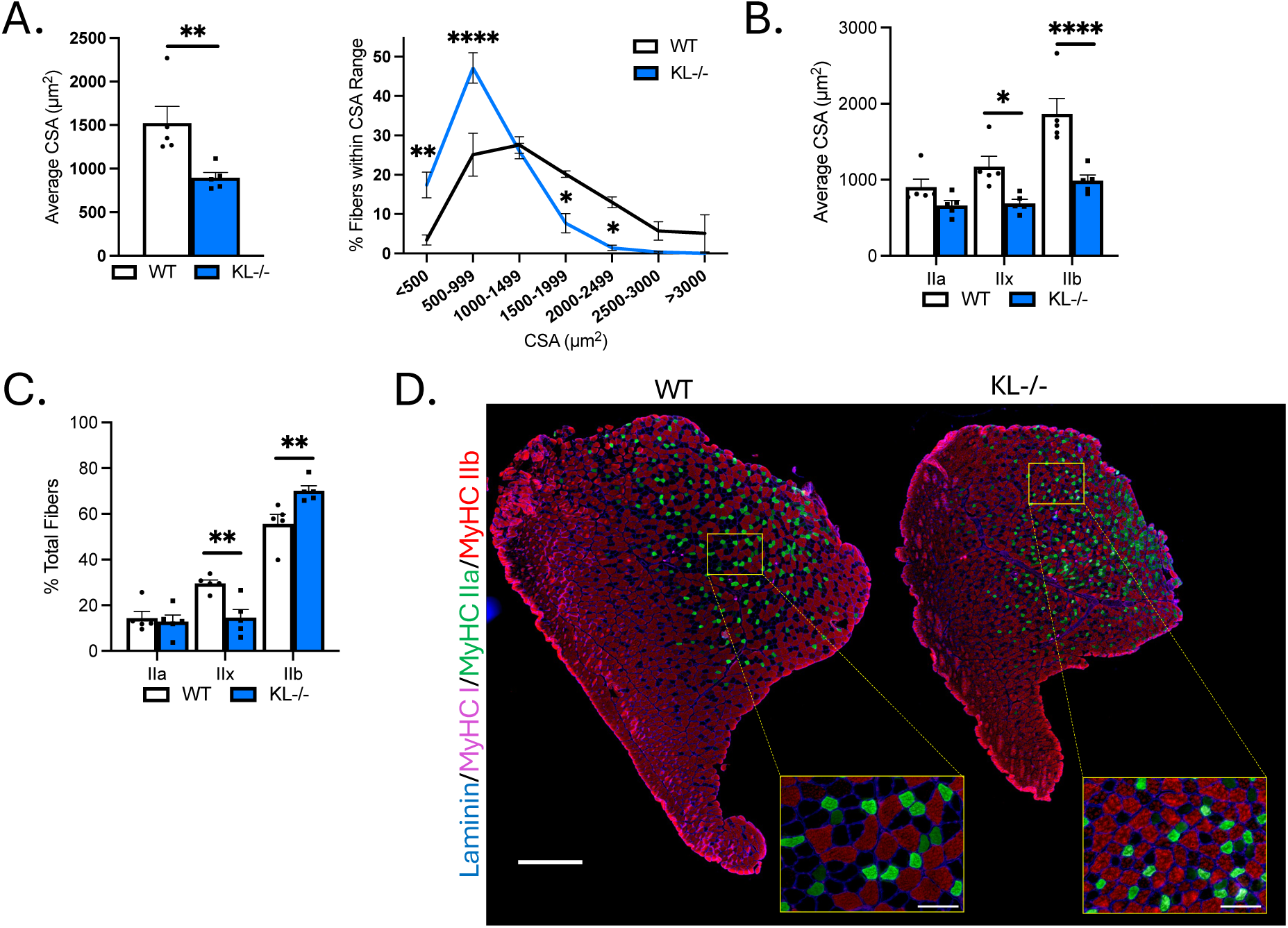
Genetic Klotho deficiency reduces muscle fiber size and affects fiber-type composition in TA muscles. (A) Left: Mean muscle fiber cross-sectional area (CSA). *P* value based on two-tailed *t* test. Right: frequency distribution of muscle fiber size by CSA. (B) Quantification of muscle fiber CSA by fiber-type in TA muscles from wild-type and KL^-/-^ mice. (C) Quantification of TA muscle composition by fiber-type. (D) Representative montages of whole TA cross-section from wild-type (left) and KL^-/-^ (right) mice immunolabeled with antibodies to laminin (blue), myosin heavy chain type (MyHC) I (magenta), MyHC IIa (green), MyHC IIb (red), and MyHC IIx unlabeled (black). Montage bar = 500 μm. Inset bar = 100 μm. N = 5 per group. Unless otherwise indicated, *P* values are based on two-way ANOVA with Sidak multiple comparison test (B-D, G-I). \**P*<0.05, \*\**P*<0.01, \*\*\*\**P*<0.00001 *versus* wild-type.

### Klotho-deficiency reduces myofiber caliber but does not affect muscle composition in slow-twitch soleus muscles

To test for differential responses of slow-twitch muscles to Klotho-deficiency, fiber-type and morphological analyses were performed on soleus. TA muscles are comprised primarily of type IIx and IIb fast/glycolytic fibers and soleus consists of mostly type I and IIa slow/oxidative fibers. Mean CSA of soleus myofibers was reduced in KL^-/-^ mice (−53%, P<0.01) associated with an increased frequency of small myofibers (<500 µm^2^, P<0.0001, Fig. 3A). CSA was reduced in types I (−42%, P<0.05), IIa (−60%, P<0.01) and IIx fibers (−57%, P<0.01, Fig. 3B, C) indicating that slow/oxidative fiber-types are susceptible to the effects Klotho-deficiency in soleus. No difference in soleus composition was observed (Fig. 3C-D). These data indicate distinct effects of Klotho-deficiency on muscle composition and type IIa myofiber size between TA and soleus muscles.

**Figure 3:**
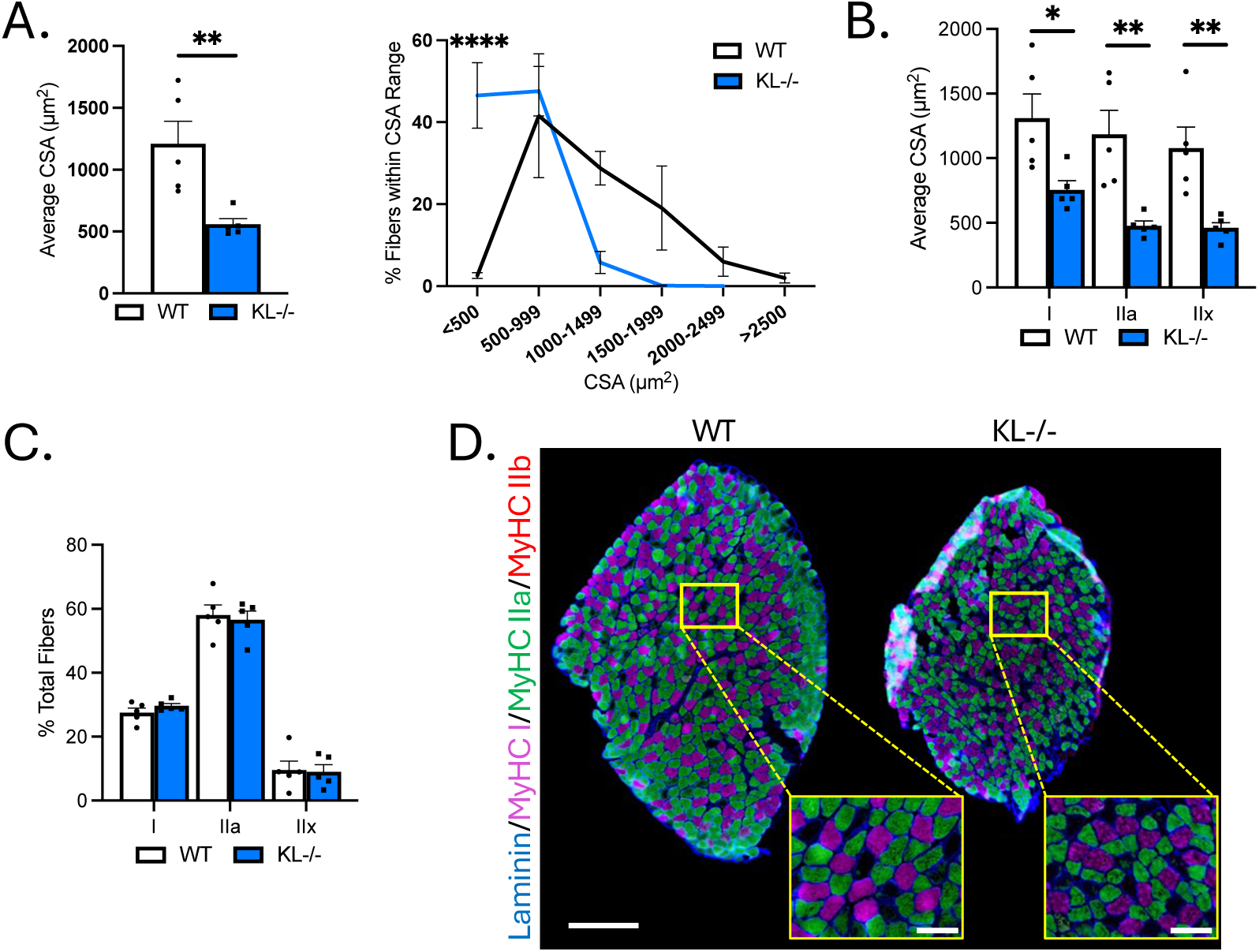
Genetic Klotho deficiency reduces muscle fiber size in soleus muscles. (A) Left: Mean muscle fiber cross-sectional area (CSA). *P* value based on two-tailed *t* test. Right: frequency distribution of muscle fiber size by CSA. (B) Quantification of muscle fiber CSA by fiber-type in soleus muscles from wild-type and KL^-/-^ mice. (C) Quantification of soleus muscle composition by fiber-type. (D) Representative montages of whole soleus muscle cross-section from wild-type (left) and KL^-/-^ (right) mice immunolabeled with antibodies to laminin (blue), myosin heavy chain type (MyHC) I (magenta), MyHC IIa (green), MyHC IIb (red), and MyHC IIx unlabeled (black). Montage bar = 250 μm. Inset bar = 50 μm. N = 5 per group. Unless otherwise indicated, *P* values are based on two-way ANOVA with Sidak multiple comparison test (B-D, G-I). \**P*<0.05, \*\**P*<0.01, \*\*\*\**P*<0.00001 *versus* wild-type.

### Genetic ablation of Klotho increases glycolytic metabolic activity in TA muscles

Because muscle fiber-type switching can affect the metabolic properties of muscle, we performed histochemical analysis on TA muscles to determine if Klotho-deficiency affects the aerobic-oxidative and anaerobic-glycolytic activities. Serial sections were used for identification of fiber-types by immunofluorescence or for histochemical analysis of SDH and GPDH activities (Figs. 4A-C). As expected, SDH and GPDH activity was highest in type IIa and IIb fibers, respectively. No change in SDH activity was detected with Klotho-deficiency (Fig. 4A). However, we observed a nearly significant (P=0.06) increase in GPDH activity of type IIb fibers and significant main effect of genotype on GPDH activity in KL^-/-^ muscles (P<0.05, Fig. 4B). SDH and GPDH activities of all fibers, independent of sub-type, was also assessed (Figs. 4D-E). No change in mean and SDH activity distribution were observed (Fig. 4D). However, consistent with a higher proportion of type IIb fibers, mean GPDH activity increased (+14%, P<0.05) and we noted a higher frequency of fibers with high GDPH activity (Fig. 4E). These findings show that changes in muscle fiber-type composition are coupled with a corresponding shift towards increased glycolytic metabolic activity in TA muscles from Klotho-deficient mice.

**Figure 4:**
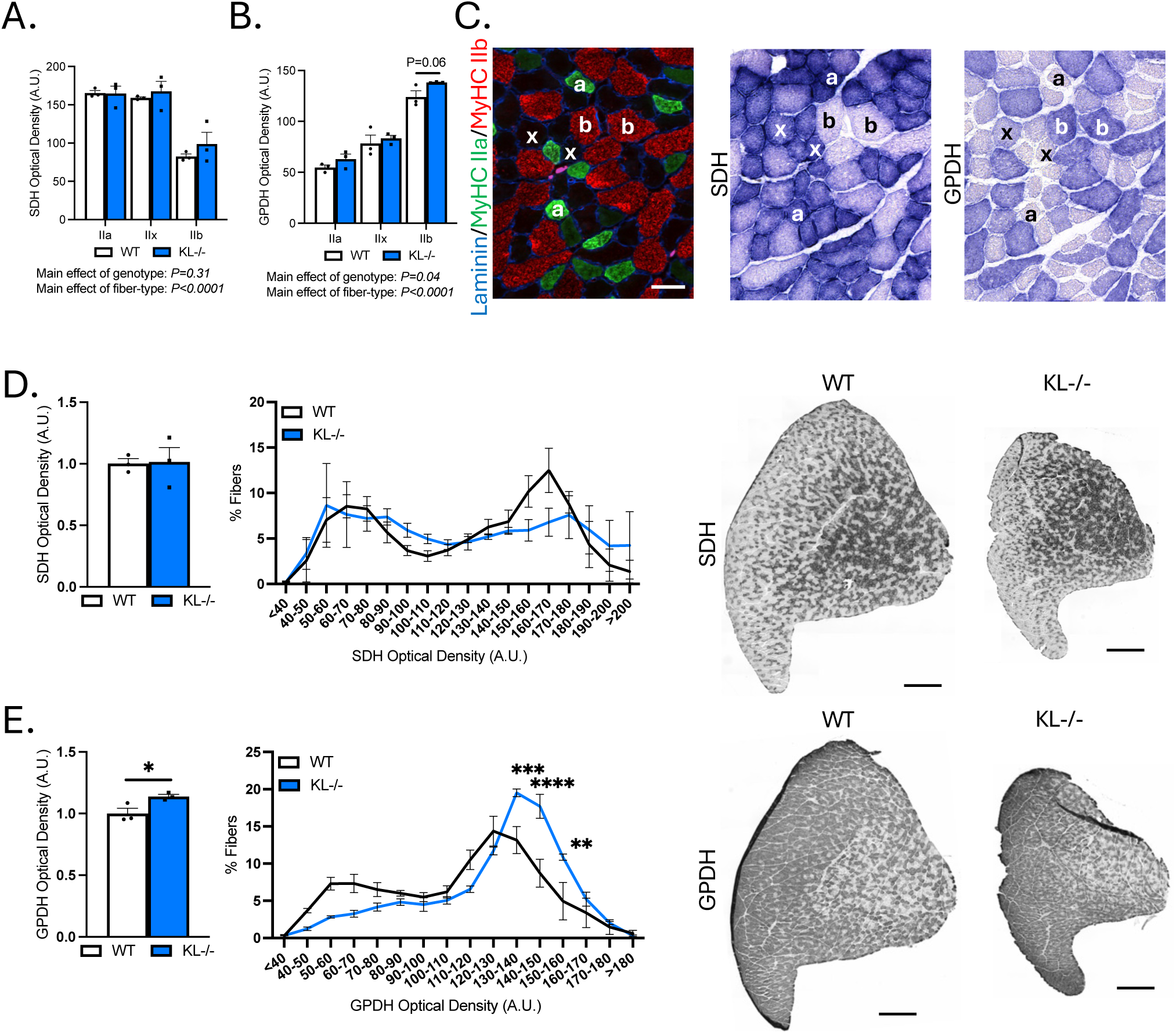
Genetic ablation of Klotho increases glycolytic GPDH enzymatic activity in TA muscles. (A-C) Serial cryosections of TA muscles immunolabeled for MyHC isoforms or succinate dehydrogenase (SDH) or glycerol-3-phosphate-dehydrogenase (GPDH) enzyme histochemistry. Quantification of SDH (A) and GPDH (B) optical density by fiber-type. *P* values based on two-way ANOVA with Tukey’s multiple comparison test. (C) Representative images of the same region of TA cross-sections. Left: Cross-sections immunolabeled for laminin (blue), MyHC1 (magenta), MyHC2a (green), MyHC2b (red), and MyHC2x unlabeled (black). Middle: SDH enzyme histochemistry. Right: GPDH enzyme histochemistry. Symbols indicate: a = type IIa, x = type IIx, and b = type IIb fiber-types across images. Bar = 50 μm (D) Left: Mean SDH enzyme activity. *P* value based on two-tailed *t* test. Middle: Frequency distribution SDH activity independent of fiber-type. Right: Representative images of SDH activity in whole TA montages from wild-type and KL^-/-^ mice in gray-scale. Montage bar = 500 μm. (E) Left: Mean GPDH enzyme activity. *P* value based on two-tailed *t* test. Middle: Frequency distribution GPDH activity independent of fiber-type. *P* values based on two-way ANOVA with Sidak multiple comparison test. Right: Representative images of GPDH enzyme histochemistry of whole TA cross-section montages from wild-type and KL^-/-^ mice in gray-scale. Montage bar = 500 μm. N = 3 per group. \**P*<0.05, \*\**P*<0.01, \*\*\**P*<0.001, \*\*\*\**P*<0.00001 *versus* wild-type.

### Increased transcriptional expression of sarcomeric and synaptic genes in Klotho-deficient muscles

To assess the molecular mechanisms behind muscle wasting, composition, and metabolic activities observed with Klotho-deficiency, we performed RNA-sequencing analysis. Multi-dimensional scaling showed distinct separation of WT and KL^-/-^ TA muscles (Fig. 5A). There were 2124 differentially expressed genes utilizing a false discovery rate <0.05 threshold, 1046 of those differed by at least log_2_0.5-fold, and of those 721 genes were upregulated and 392 downregulated (Fig. 5B). Candidate genes were examined for expressional changes (Fig. 5C). We observed increased expression of Mstn, a negative regulator of muscle growth, but no change in the expression of pro-atrophy genes Fbxo32 or Trim63. The expression of sarcomeric transcripts Myh3, Myh4, Myh6, Myh7b, Myh8, Myh13, Myl4, Myl10, Tnnc1, Tnni1, and Tnnt2 were elevated in KL^-/-^ muscle.

**Figure 5:**
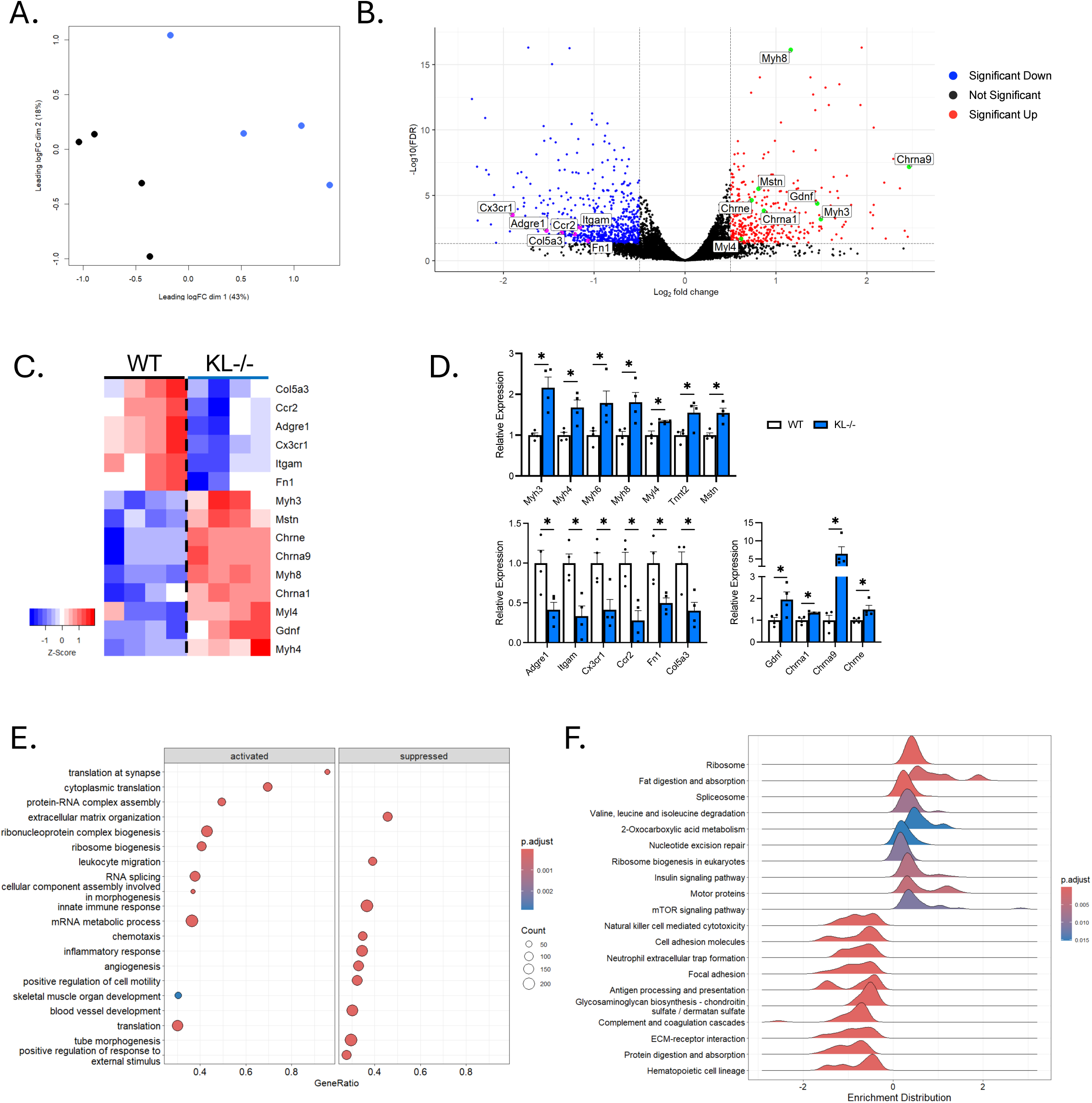
RNA-sequencing analysis shows distinct gene expression profiles in skeletal muscles from Klotho-deficient mice. (A) Multi-dimensional scaling plot comparing TA muscles from wild-type and KL^-/-^ mice. (B) A volcano plot highlighting selected differentially expressed genes (DEGs). Increased expression (red) and decreased expression (blue) in KL^-/-^ muscles. Colored dots = abs(log_2_0.5) fold change with FDR<0.05. (C) A heatmap of samples showing select DEGs in WT and KL^-/-^ muscles. Positive (red) and negative (blue) Z-score. (D) qRT-PCR assays validating differential expression of selected sarcomeric (top), inflammatory and pro-fibrotic (bottom left), and acetylcholine receptor and neurotrophic genes (bottom right). (E) Dot plot depicting activated (left) and suppressed (right) GO of Biological Process terms in KL^-/-^ muscles. (F) Ridgeline plot, grouped by gene set, representing activated and suppressed KEGG pathways in KL^-/-^ muscles. N = 4 per group.

Increased Myh4 expression was consistent with a higher concentration of type IIb myofibers (Fig. 2C). Myh3, Myh8, Myl4, and Tnnt2 genes are expressed highest during development, their expression is downregulated postnatally but re-expressed in response to muscle injury or denervation (Sartore *et al*., 1982; Schiaffino *et al*., 1988; Saggin *et al*., 1990). In this regard, transcripts for acetylcholine receptors Chrna1, Chrna9, and Chrne and motor neuron survival factor Gdnf were elevated in KL^-/-^ muscle. However, we did not observe transcriptional changes associated with muscle damage. No change in expression of myogenic factors Pax7, Myod1, Myog, Myf5, and Myf6, and reduced expression of inflammatory and extracellular remodeling genes, including macrophage markers Adgre1 and Itgam, chemokine receptors Cx3cr1 and Ccr2, and extracellular matrix components Fn1 and Col5a3. Expressional changes of key transcripts were validated by qRT-PCR (Fig. 5D).

To identify biological processes in muscle affected by Klotho-deficiency, GSEA and GO analyses were performed. We found positive enrichment for translation at the synapse, cytoplasmic translation, protein-RNA complex assembly, ribosome biogenesis, and skeletal muscle development in KL^-/-^ muscle (Fig. 5E). In contrast, extracellular matrix organization, leukocyte migration, and innate immune response were downregulated terms. KEGG pathway enrichment analysis was also performed to gain mechanistic insights (Fig. 5F), top enriched pathways include ribosome and spliceosome processes which play essential roles in normal cell physiology regulating protein synthesis and gene expression. Other enriched pathways include increased valine, leucine, and isoleucine degradation and 2-oxocarboxylic acid metabolism. In accord with Klotho repressing insulin and IGF-1 signaling (Kurosu *et al*., 2005), enrichment of insulin signaling and mTOR signaling were observed in KL^-/-^ muscles. Interestingly, mTOR signaling is a primary driver of muscle wasting and NMJ dysfunction (Castets *et al*., 2013; Ham *et al*., 2020). Collectively, these data raise the possibility that Klotho-deficiency facilitates changes to the muscle synapse promoting muscle wasting and altered expression of sarcomeric transcripts.

### Klotho-deficiency promotes muscle weakness

The observations that Klotho-deficiency affects myofiber size and composition indicates that Klotho may affect muscle function. We tested that possibility by performing *in vivo* plantarflexion functional analysis. Absolute muscle twitch was 48% lower indicating reduced force to a single impulse in KL^-/-^ mice (P<0.01, Fig. 6A). However, there was no effect on muscle twitch scaled to body weight (Fig. 6A). Muscle twitch maximal rate of contraction (−54%, P<0.001) and relaxation (−61%, P<0.001, Fig. 6A) were reduced in KL^-/-^ mice.

**Figure 6:**
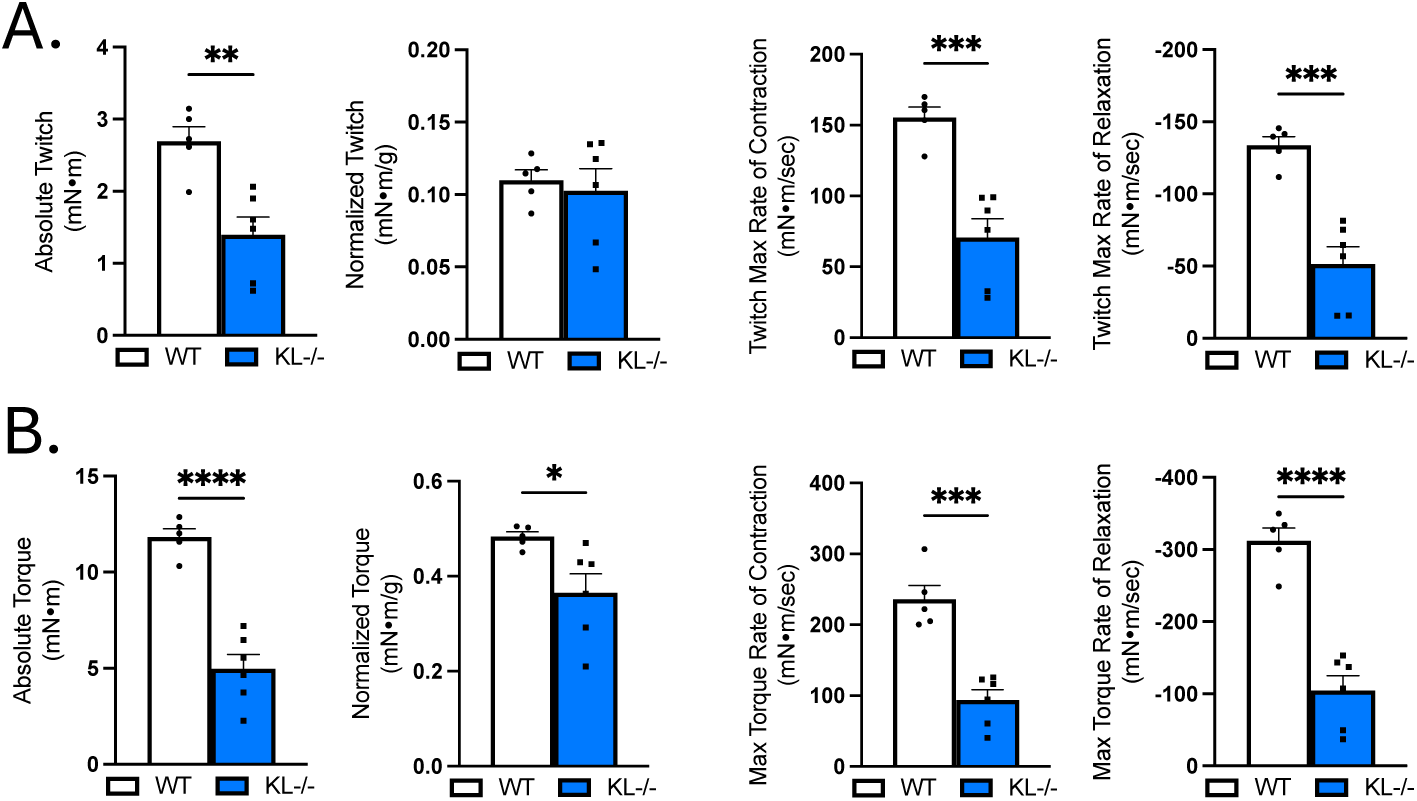
Muscle contractile function is impaired in Klotho-deficient mice. *In vivo* plantar flexion force assessment of the triceps surae muscle group in WT and KL^-/-^ mice reported as: (A) absolute twitch (mN·m), twitch normalized to body weight (mN·m per g body weight), twitch max rate of contraction and relaxation; (B) absolute torque (mN·m), tetanic torque normalized to body weight (mN·m per g body weight), max torque rate of contraction and relaxation. N = 5-6 per group. All *P* values are based on two-tailed *t* test. \**P*<0.05, \*\**P*<0.01, \*\*\**P*<0.001, \*\*\*\**P*<0.0001

Reductions in absolute torque (−58%, P<0.0001) and tetanic torque normalized to body weight (−23%, P<0.05, Fig. 6B) indicates muscle weakness independent of differences in body mass in KL^-/-^ mice. Maximal rate of torque development (−60%, P<0.001) and relaxation (−66%, P<0.0001, Fig. 6B) also decreased in KL^-/-^ mice. Taken together, Klotho-deficiency promotes muscle weakness and alters muscle contraction and relaxation rates.

### Motor unit connectivity is impaired in Klotho-deficient mice

Given that Klotho-deficiency promotes weakness, alters contraction and relaxation dynamics, and transcriptional analysis shows changes associated to the muscle synapse, we next sought to test indices of motor unit connectivity. Electrophysiological measurements showed no change in baseline-to-peak assessment of CMAP (Fig. 7A). However, SMUP increased 3.6-fold in KL^-/-^ mice relative to WT (P<0.01, Fig. 7B).

**Figure 7:**
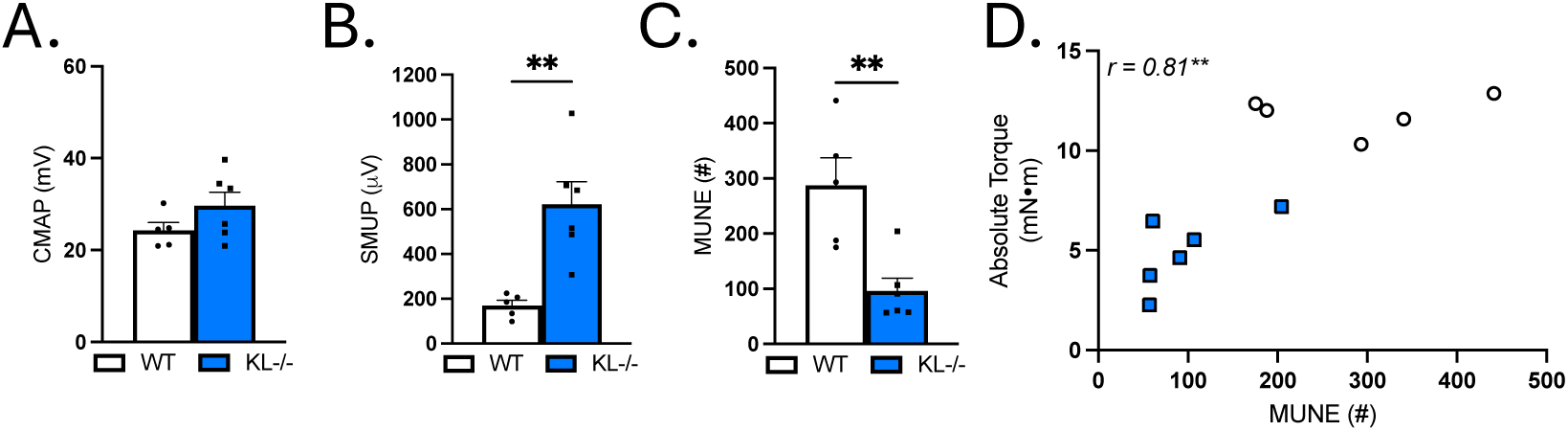
Genetic Klotho-deficiency alters single motor unit potential (SMUP) and motor unit number estimation (MUNE). (A) Compound muscle action potential (CMAP; millivolts (mV)). (B) SMUP (microvolts (μV)). (C) MUNE (number of motor units (#). (D) *In vivo* plantar flexion absolute torque correlated with MUNE. N = 5-6 per group. All *P* values are based on two-tailed *t* test (A-C) or correlation analysis for Pearson r value. \*\**P*<0.01.

MUNE calculations showed a robust 67% reduction indicating reduced motor units with Klotho-deficiency (P<0.01, Fig. 7C). Correlation analysis revealed that MUNE positively associated with absolute torque (Pearson r=0.81, P<0.01, Fig. 7D). These data indicate that impaired motor unit connectivity is associated with muscle weakness in Klotho-deficient mice.

### Klotho-deficiency alters the structure and synaptic overlap at the NMJ

Muscle wasting, weakness, altered composition, and functional loss in motor unit innervation could result from changes to the NMJ. First, we quantified the number of myonuclei per fiber to test for effects on myonuclear accretion. No change indicates that reduced myofiber size was not associated with effects to the myonuclear domain (Fig. 8A). To address effects of Klotho-deficiency on muscle injury, serum CK was analyzed and the proportion of centrally nucleated fibers (CNFs) were quantified. Serum CK levels did not change with Klotho-deficiency (Fig. 8B). However, there was a small but significant increase in CNFs (WT: 0.2% *vs.* KL^-/-^: 0.7%, P < 0.05, Fig. 8C). Central nucleation is a well-recognized indicator of regeneration but changes in nuclear placement may also occur with denervation(Schmalbruch, 1976; Borisov *et al*., 2001). To address this point, we assessed myofiber expression of NCAM whose expression accumulates intracellularly in denervated fibers (Covault & Sanes, 1985). Indeed, the percentage of NCAM+ fibers increased in KL^-/-^ muscles. NCAM was most prevalent in small, angulated fibers consistent with denervation (P<0.0001, Fig. 8D).

**Figure 8:**
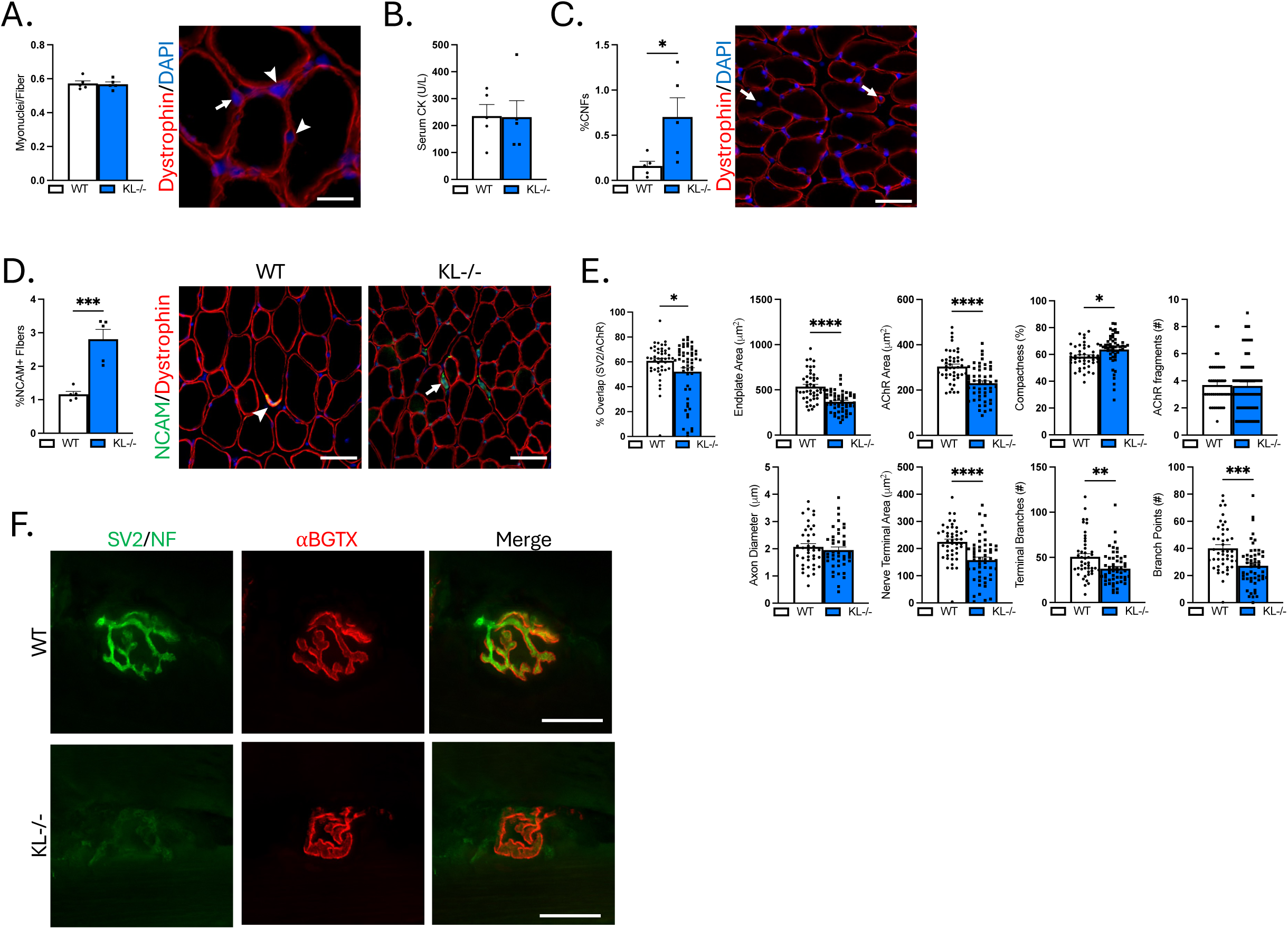
Genetic ablation of Klotho increases concentration of denervated muscle fibers, reduced synaptic contact, and alters NMJ morphology. (A) Myonuclei number per muscle fiber. N=5/group. Representative image of muscle cross-section immunolabeled with dystrophin and stained with DAPI to visualize nuclei for quantification of myonuclei. Arrow = myonuclei. Arrowhead = interstitial nuclei. Bar = 20 μm. (B) Serum creatine kinase (CK) levels. N=5/group. (C) Quantification of the percentage of muscle fibers with centrally located nuclei to total muscle fibers. N=5/group. Representative image of muscle cross-section immunolabeled with dystrophin and stained with DAPI to quantify centrally nucleated fibers. Arrow = centrally nucleated myofiber. Bar = 50 μm. (D) Quantification of the proportion of denervated muscle fibers expressing NCAM. N=5/group. Representative images of muscle sections immunolabeled with dystrophin (red) and NCAM (green) from wild-type (left) and KL^-/-^ mice (right). Arrowhead = NMJ with high expression of NCAM. Arrow = small angulated denervated fiber with high cytosolic expression of NCAM. Bar = 50 μm. (E) Quantification of NMJ morphological properties of longitudinal sections of TA muscles from wild-type and and KL^-/-^ mice. Percent co-localization of synaptic vesicles and AChRs; post-synaptic NMJ morphology endplate area, AChR area, compactness (AChR area/endplate area), and number of AChR fragments; and pre-synaptic NMJ morphology axon diameter, nerve terminal area, number of terminal branches, and branch points. In dispersion plots, each point represents a single NMJ from n=5 wild-type and n=6 KL^-/-^ biological replicates, n=6-15 en-face NMJs quantified per biological replicate. F) Representative images of longitudinally cut TA muscle sections from wild-type (top) and KL^-/-^ mice (bottom) stained with bungarotoxin (red) and immunolabeled with SV2 and neurofilament (green). Bar = 20 μm. All *P* values are based on two-tailed *t* test. \**P*<0.05, \*\**P*<0.01, \*\*\**P*<0.001, \*\*\*\**P*<0.0001.

To gain insight into whether Klotho-deficiency affects NMJ structure, morphological analysis was performed on longitudinal sections stained with antibodies to neurofilament to identify the axon, SV2 to label the nerve terminal, and ⍺-bungarotoxin to visualize post-synaptic AChRs (Fig. 8E-F). Overlap between pre-synaptic and post-synaptic components decreased indicating reduced area of synaptic contact in KL^-/-^ mice (−14%, P<0.05, Fig. 8E). Pre- and post-synaptic morphology were also altered with Klotho-deficiency (Fig. 8F). Endplate area (−32%, P<0.0001) and AChR area (−24%, P<0.0001) decreased, whereas compactness of AChRs at the endplate increased (+9%, P<0.05, Fig. 8E). At the pre-synapse, nerve terminal area (−30%, P<0.0001), number of terminal branches (−26%, P<0.01) and branch points (−32%, P<0.001, Fig. 8E) decreased in KL^-/-^ mice. Taken together, these findings demonstrate that the functional loss of motor unit connectivity is associated with increased prevalence of NCAM+ denervated fibers, reduced area of synaptic contact, and morphological changes to the NMJ potentially affecting signal transmission and muscle function.

## DISCUSSION

The major findings of the current study are that Klotho affects muscle mass, composition, function, and motor unit connectivity. These conclusions are based on the following experimentation using Klotho-deficient mice. First, we demonstrated reductions in body weight, lean mass, muscle mass, and myofiber caliber. Second, we showed altered muscle composition with an increased proportion of type IIb fibers and elevated glycolytic GPDH activity in KL^-/-^ muscles. Transcriptomic data revealed positive enrichment for biological processes related to translation at the synapse, as well as increased expression of genes encoding AChRs and sarcomeric genes that are re-expressed in mature muscles after denervation. Finally, we showed that transcriptional changes were coupled with functional loss of motor unit number, increased prevalence of denervated fibers, reduced area of synaptic contact, and small, compact NMJ morphology. Decreased muscle torque also correlated with reductions in motor unit number.

We were very interested to learn that Klotho-deficiency resulted in transcriptional, morphological, and functional changes impacting motor unit connectivity. To our knowledge, this is the first report to demonstrate changes to the NMJ and muscle denervation related to Klotho-deficiency. However, excess Klotho promotes motor neuron survival (Zeldich *et al*., 2019). Altered motor nerve function, muscle loss, and weakness are common pathologies observed in clinical conditions of Klotho insufficiency (Tomlinson & Irving, 1977; Krishnan *et al*., 2005; Kaya *et al*., 2013). Functional consequences related to the loss of neural input in muscle include diminished absolute torque, lower rates of force development and relaxation, and fatigue that manifest as a reduction in strength and coordination resulting in increased susceptibility to falls (Power *et al*., 2013).

Taken together, our findings support that changes to the muscle synapse affecting motor unit connectivity and muscle function may contribute to increased risk of falls and frailty observed in clinical populations with low Klotho levels (Shardell *et al*., 2019; Sanz *et al*., 2021; Veronesi *et al*., 2021).

We also observed large reductions in muscle mass and myofiber caliber with Klotho-deficiency. Muscles are comprised of fibers classified by sub-type based on fatigability, contractile and metabolic properties.

Considering that different muscles and fiber-types within a muscle may respond differently to the same stimulus, we were interested to learn that TA muscles of Klotho-deficient mice had smaller type IIx and IIb but not type IIa fibers. In contrast, all fiber-types were smaller in soleus. Like our observations, type IIa fibers of the TA are resistant to denervation-induced atrophy with neurodegenerative disease (Villalón *et al*., 2019).

However, in denervated soleus muscles both type I and type II fibers atrophy (Jaweed *et al*., 1975).

Another intriguing observation is that Klotho-deficiency altered composition of the TA, but not soleus. TA muscles had a higher percentage of type IIb fibers to the detriment of type IIx fibers. Fiber-type is a crucial determinant of contractile function and metabolism. Type IIb fibers rely on glycolysis for energy production.

Consistent with this, GPDH enzymatic activity increased in TA muscles from Klotho-deficient mice. Neural input directly influences fiber-type, a slow-to-fast fiber-type switch occurs when slow fibers are re-innervated by fast motor neurons (Buller *et al*., 1960).

The mechanisms underlying muscle wasting in Klotho-deficient mice are multiple. Klotho affects several signaling pathways known to regulate muscle mass (Ohsawa *et al*., 2023; Heitman *et al*., 2024; Bose *et al*., 2025), we used RNA-sequencing for unbiased characterization of transcriptional changes to muscles.

Surprisingly, expression of key regulators of muscle atrophy Fbxo32 and Trim63 were not differentially expressed but Mstn expression was elevated. GO analysis showed activation of pathways opposed to catabolism, including protein translation and ribosome biogenesis. One potential explanation for this is that protein synthesis rates of individual fibers increase to mitigate atrophy after denervation (You *et al*., 2021). In our model, denervation occurs spontaneously likely complicating our interpretation. Indeed, others reported increased expression of Fbxo32 and Trim63 in gastrocnemius of Klotho-deficient mice (Heitman *et al*., 2024). Provided that we also observe smaller myofibers a negative net balance in protein turnover is obvious.

We did not observe an impaired myogenic response in normal muscles of Klotho-deficient mice. Muscle progenitor cells play a crucial role in postnatal growth fusing to fibers to add nuclei to accommodate increased synthetic demand (Bachman *et al*., 2018). In our study, we found no change to the number of myonuclei per fiber. Therefore, muscle wasting does not appear to be related to differences in myonuclear accretion. A role for Klotho in the regulation of muscle progenitor cell function affecting regeneration in response to injury is well established (Wehling-Henricks *et al*., 2016; Ahrens *et al*., 2018; Sahu *et al*., 2018; Wehling-Henricks *et al*., 2018; Welc *et al*., 2020). However, serum CK levels were normal, fewer than 1% of myofibers contained central nuclei, and transcriptional expression of key regulators of myogenesis were normal in Klotho-deficient mice. Collectively, these data suggest reductions in muscle mass are not related to impaired myogenesis or injury. An important next step in future investigations will be to use models for the inducible-deletion of Klotho to further distinguish effects of Klotho-deficiency on adult, mature muscles.

The specific mechanisms through which Klotho-deficiency alters the structure and function of the NMJ affecting innervation and wasting are not yet known. We used RNA-sequencing and pathway analysis to attempt to identify a mechanism through which Klotho-deficiency affects muscle innervation. Although bioinformatic inquiry cannot prove a causal mechanism, the data identified activation of insulin and mTOR signaling pathways and activation of associated biological processes downstream, like ribosome biogenesis. Klotho mutant mice have increased insulin sensitivity (Utsugi *et al*., 2000). mTOR coordinates cellular proliferation, growth, and survival with environmental factors, including nutrient availability and growth factors controlling muscle mass (Yoon, 2017). Furthermore, insulin and IGF-1 signaling enhances activation of mTOR signaling (Sarbassov *et al*., 2005). Emerging evidence supports a key role for mTOR in regulating maintenance of the post-synaptic endplate to preserve NMJ integrity (Castets *et al*., 2019; Ham *et al*., 2020; Ang *et al*., 2022). Loss of motor unit connectivity may precede declines in muscle mass and contractility and therefore causally contribute to muscle wasting and weakness.(Sheth *et al*., 2018) This indicates that aberrant activation of insulin and mTOR signaling pathways with Klotho-deficiency may alter NMJ morphology reducing motor unit connectivity affecting muscle mass and function.

Although our findings show profound effects of Klotho-deficiency on skeletal muscle, we are unable to conclude the contribution of muscle Klotho expression to these effects. The function of skeletal muscle Klotho expression is largely unknown. Here, we confirmed ∼130 kDa Klotho protein expression in muscle homogenates. Prior works showed that transcriptional expression of Klotho in muscles is suppressed transiently after injury (Welc *et al*., 2020), with aging (Wehling-Henricks *et al*., 2018), and after disease onset in Duchenne muscular dystrophy, a muscle wasting disorder (Wehling-Henricks *et al*., 2016). In support of a role for muscle-derived Klotho, muscle targeted knockdown by RNA interference was shown to inhibit myogenesis impairing regeneration after injury (Sahu *et al*., 2018). Whereas other works attribute effects of

Klotho-deficiency to in-direct effects driven by phosphate retention and iron accumulation (Heitman *et al*., 2024; Bose *et al*., 2025). Also plausible are combinatorial effects through which reduced muscle Klotho expression impacts the muscle’s capacity to respond to endocrine or metabolic stress. To this end, in the absence of the influence of endocrine factors, knockdown of Klotho in normal primary muscle cell cultures *in vitro* induced cellular senescence, mitochondria dysfunction and altered bioenergetics (Sahu *et al*., 2018). Muscle cells that acquire a senescent phenotype accelerate wasting and dysfunction (Falvino *et al*., 2024).

In summary, clinical observations suggest that reduced circulating Klotho levels correlate with muscle weakness, frailty, increased risk of falls, and reduced ability to perform activities of daily living. Therefore, Klotho may play an important role in the pathogenesis of muscle wasting and weakness. Our findings identify a previously unrecognized role for Klotho in regulating the structure and function of the NMJ affecting motor unit connectivity. Mechanistically, our data support that Klotho-deficiency causes muscle wasting and weakness at least in part due to loss of neuromuscular integrity. Supporting this conclusion is a strong correlation between motor unit number and muscle torque. Together, these results advance our understanding of relationship between Klotho, muscle wasting and weakness.

## ACKNOWLEDGEMENTS

Research reported in this publication was supported by the National Heart Lung and Blood Institute under award numbers R01HL158647 (SSW) and the National Institute of Arthritis Musculoskeletal and Skin Disease T32AR065971 (ASJ). Additional support was provided by Muscular Dystrophy Association #603201 (SSW), Indiana University School of Medicine and Indiana Center for Musculoskeletal Health as new laboratory start-up funds (SSW). Sequencing analysis was carried out in the Center for Medical Genomics at Indiana University School of Medicine, which is partially supported by the Indiana University Grand Challenges Precision Health Initiative. The #BA-D5, #SC-71, #BF-F3 (developed by Schiaffino, S.), #Mandys8(8H11) (developed by Morris, G.E.), #SV2 (developed by Buckley, K.M.), and #2H3 (developed by Jessell, T.M. & Dodd, J.) antibodies were obtained from the Developmental Studies Hybridoma Bank, created by the NICHD of the NIH and maintained at The University of Iowa, Department of Biology, Iowa City, IA 52242

## CONTRIBUTIONS

Study conceptualization and design, L.A.B., C.T., J.F.V., K.E.W., J.R.H., and S.S.W.; methodology, acquisition of data, and/or interpretation of data, L.A.B., C.T., J.F.V., A.J.F., A.J.S., A.A., H.W., G.D.S., J.R.H., and S.S.W.; formal analysis, L.A.B., C.T., J.F.V., A.J.F., A.J.S., A.A., H.W., G.D.S., J.R.H., and S.S.W.; writing—original draft preparation, S.S.W.; writing—review and editing, L.A.B., C.T., J.F.V., A.J.F., A.J.S., A.A., H.W., G.D.S., K.E.W., J.R.H., and S.S.W.; funding acquisition, S.S.W.. All authors have read and approved the final version of the manuscript. All authors have agreed to be personally accountable for their contributions to ensure that questions related to the accuracy or integrity of any part of the work are appropriately investigated, resolved, and the resolution documented in the literature.

